# Detection of West Nile Virus via Retrospective Mosquito Arbovirus Surveillance in the United Kingdom

**DOI:** 10.1101/2025.06.17.659932

**Authors:** Robert C. Bruce, Anthony J. Abbott, Ben P. Jones, Bathsheba L. Gardner, Estela Gonzalez, Andra-Maria Ionescu, Madhujot Jagdev, Ava Jenkins, Mariana Santos, Katharina Seilern-Macpherson, Hugh J. Hanmer, Robert A. Robinson, Alexander G.C. Vaux, Nicholas Johnson, Andrew A. Cunningham, Becki Lawson, Jolyon M. Medlock, Arran J. Folly

## Abstract

In March 2025, as part of on-going enhanced surveillance for mosquito-borne *Orthoflaviviruses*, West Nile virus (WNV) RNA was detected in two pools of female *Aedes vexans* collected in July 2023 in Nottinghamshire, England. Viral RNA detection was achieved via reverse transcription-qPCR. Sequencing and subsequent phylogenetic analysis of a 402 bp fragment indicated that the detections clustered with WNV lineage 1a. The exact origin of this virus remains unclear, but this finding indicates a historic WNV presence in the United Kingdom (UK). Surveillance has not provided evidence of further viral transmission to date.

---

West Nile virus is a single-stranded RNA virus that is maintained in an enzootic cycle between mosquitoes and wild birds [1]. The virus has been detected in mainland Europe since the mid-1990s [2] where two lineages (1a and 2) circulate and cause sporadic outbreaks, with disease recorded in wild birds [3,4], horses [5] and humans [6,7]. The virus has steadily expanded across Western Europe, including recent detections in Germany and the Netherlands [8,9], but so far with no evidence of emergence in the UK. Passive surveillance of dead wild birds in the UK for *Orthoflaviviruses* has not detected WNV RNA in any of the 5,039 individuals sampled 2013-2024, inclusive [10].

Following the emergence and establishment of Usutu virus (USUV, *Orthoflavivirus*) in the UK in 2020 [11], enhanced arbovirus surveillance under the One Health paradigm was established through the VB-RADAR (Vector-borne Real-time Arbovirus Detection And Response) project (https://www.vb-radar.com). This project included screening of mosquitoes from a range of wetland and urban areas deemed at high risk of arboviral introduction [12]. From March 2023-March 2025, inclusive, 31,582 mosquitoes comprising 22 species (Supplementary Table 1) were collected using a range of active trapping methods at 26 sites (Figure 1) and were analysed for arbovirus infection. Two *Aedes vexans* pools tested positive for WNV RNA, with the remainder being negative. *Aedes vexans* is an uncommon species in the UK, requiring summer flooded wetlands to trigger egg hatching and larval development, and so far it is only known to occur in a few highly localised areas of the UK [13,14]. Although not considered a principal enzootic WNV vector in Europe, in contrast to *Culex pipiens s*.*s/Cx. torrentium*. and *Cx. modestus, Ae. vexans* is a potential bridge vector, as it feeds on birds, horses and humans [15,16], and is a competent vector of a number of arboviruses including WNV [17–21]. Consequently, *Ae. vexans* may play an important role in arboviral circulation, particularly where it is highly abundant, and therefore should be considered for surveillance efforts to appraise risks to public health, particularly where it is highly abundant and causes nuisance biting.

**Figure 1.**
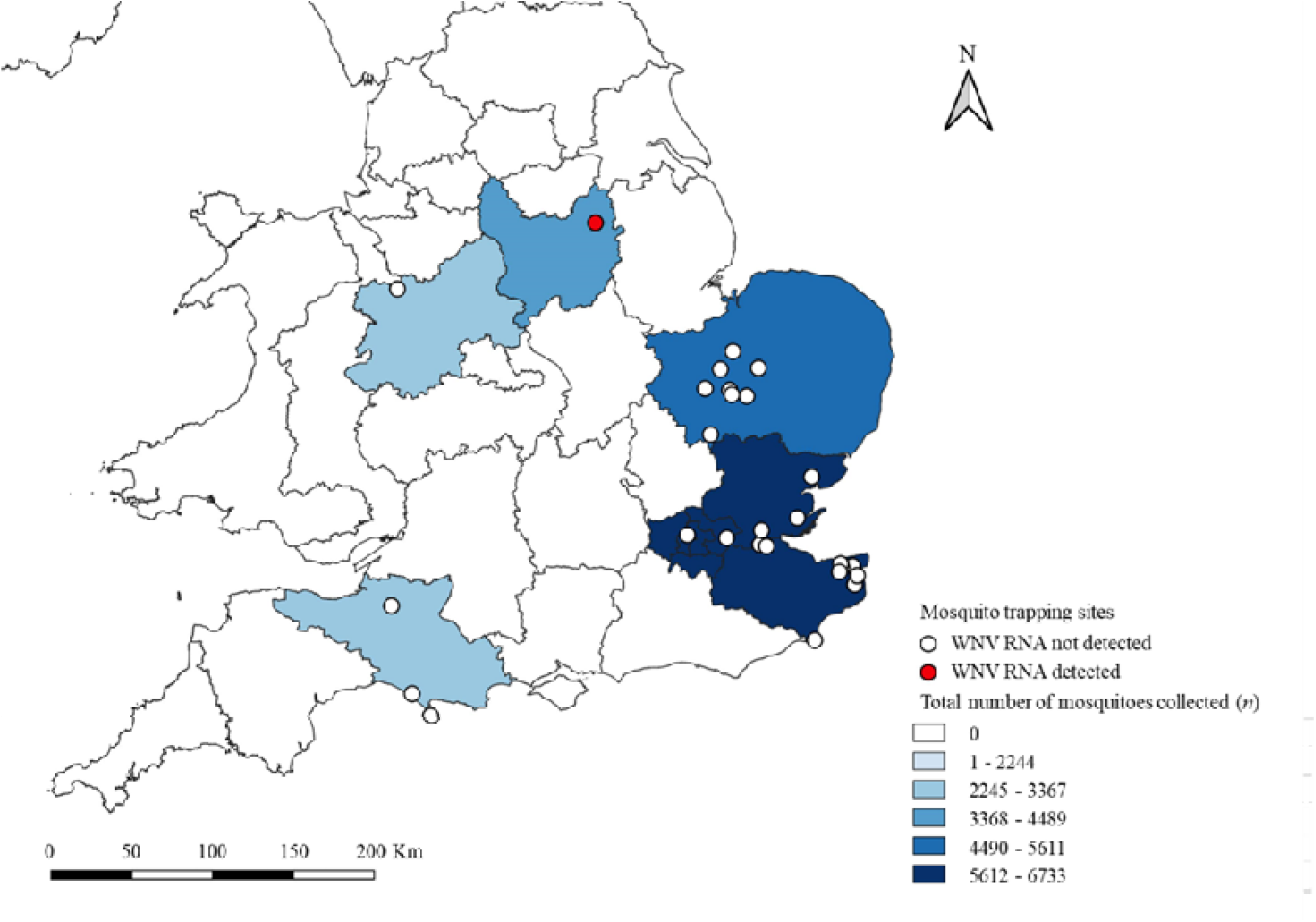
Vector-borne RADAR mosquito surveillance sites, for arbovirus screening across, March 2023-March 2025 inclusive, indicating sites, numbers collected and locations where West Nile virus was either detected or was not detected (produced using QGIS Maidenhead v3.36.1)

## Surveillance of *Aedes vexans* in Nottinghamshire and WNV detection

As part of long-running mosquito surveillance (since 2018) in Gamston (near Retford), Nottinghamshire (England) [14,22], mosquitoes were collected using a Mosquito Magnet® Executive model (Woodstream Corporation, St. Joseph, MO, USA) baited with an octenol lure. For those screened for arboviruses, traps were deployed from July–September 2023 and run every other week for four nights. A total of 44,080 female mosquitoes was caught in the 2023 trapping season. Total catch was dominated by *Ae. vexans* (99.99%), and no *Cx. Pipiens s*.*l*. or *Cx. modestus* were captured. All mosquitoes were morphologically identified using a stereomicroscope [33]. Due to laboratory capacity constraints a subset of 2,000 *Ae. vexans* individuals trapped during the period 11^th^-21^st^ July 2023 was pooled by date of capture into groups of ten prior to RNA extraction (see Supplementary Information for details regarding mosquito sampling); which was conducted using a KingFisher™ Flex Purification System (Thermo Fisher, UK).WNV-PCR-positive pools were re-extracted using a QIAamp Viral RNA Extraction mini kit (QIAGEN, UK) (Supplementary Information).

Nucleotide extracts from all 200 pools of *Ae. vexans* were screened for the presence of WNV RNA, using three PCR assays targeting the 5’-UTR and non-structural gene 5 (NS5) [23–25]. Of these, two *Ae. vexans* pools (hereby referred to as Av_1 and Av_2 [both collected on 21^st^ July 2023]) tested positive with *Ct* values ≤ 33.4. Attempts to isolate the virus in Vero cells from the original mosquito homogenate were unsuccessful. The amplicons from positive pools were submitted for Sanger sequencing to confirm their identity. Sequencing produced two amplicons matching the 5’-UTR region and NS5 of WNV lineage 1a (BLAST identity > 95%).

In an attempt to generate additional sequence data, amplicon-based sequencing was undertaken using a GridION (Oxford Nanopore Technologies, UK). Here, a multiplex PCR designed to amplify 37 consecutive sequences of the WNV lineage 1a genome was used (Supplementary Information). The GridION produced approximately 140,000 reads for Av_1 and Av_2 combined, over a fifteen-hour sequencing window, all of which were of similar size to the desired 400 bp contigs. Following quality checks and primer removal, no reads mapped to WNV lineage 1a or 2 genomes using a combination of BWA (v0.7.13) [26] and SAMtools (v1.21) [27,28]. The resulting GridION contig file was converted to a BLAST database and a representative WNV genome (GenBank accession number: OM302321) was used to identify potential matches. Five-hundred and twenty contigs from Av_1 matched WNV genome (C-terminal of the E gene and N-terminal of NS1); however, no reads from Av_2 matched the WNV template genome. We aligned 52 comparable sequences from Av_1 against a WNV genome in MEGA11 (v11.0.11) to generate a 402 bp consensus sequence for downstream phylogenetic analysis.

The *Ae. vexans* derived WNV consensus sequence was aligned with 65 WNV genomes from GenBank (Supplementary Table 2) and imported into BEAST (v1.10.4), a Bayesian phylogenetic tree was produced using the GTR+I+G4 nucleotide substitution model and 30,000,000 Markov chain Monte Carlo generations. The resulting tree was visualised and annotated in FigTree v1.4.4 (Figure 2).

**Figure 2.**
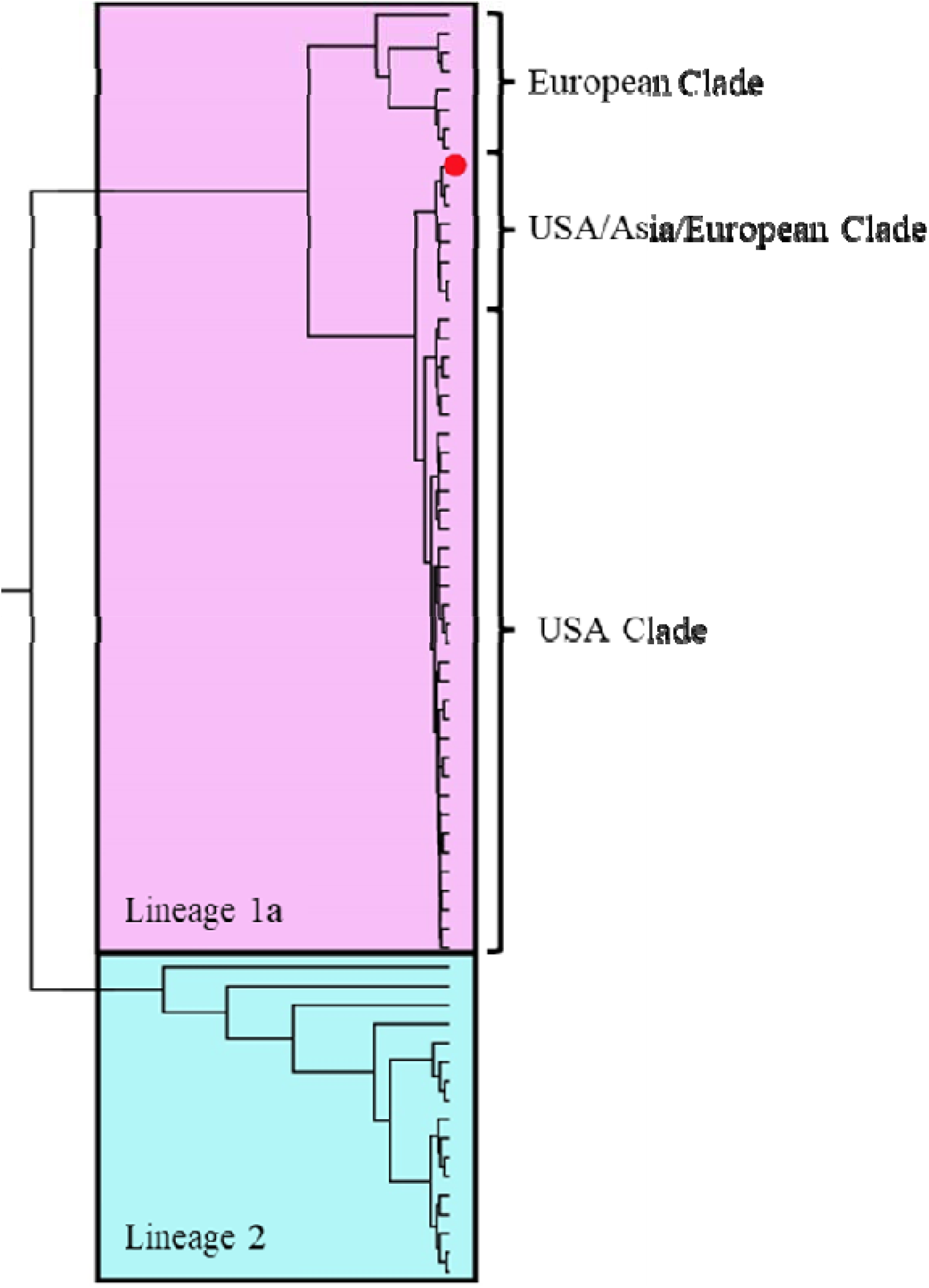
Bayesian phylogenetic tree of a 402 bp UK West Nile virus (WNV) consensus contig, obtained through Oxford-Nanopore sequencing of infected *Aedes vexans* aligned against 65 WNV genomes from GenBank. Constructed using BEAST (v1.10.4), the Bayesian phylogenetic tree was produced using the GTR+I+G4 nucleotide substitution model and 30,000,000 Markov chain Monte Carlo generations, visualised and annotated in FigTree v1.4.4 The red circle marks the UK sequence (Genbank PV659664), which clusters within WNV lineage 1a clade.

## Discussion

We present data to support a detection of WNV RNA in UK mosquitoes collected in 2023 from Nottinghamshire, England. Our results show that the detection most likely represents WNV lineage 1a, which circulates in mainland Europe and the USA, having a large impact on native bird populations [29,30]. Despite ongoing surveillance in wild birds and mosquitoes at a larger number of field sites across England, we have no evidence of further viral circulation in the UK to date.

*Aedes vexans* is an opportunistic feeder which can occur in locally high densities, and therefore may have a role in zoonotic arbovirus transmission, including WNV [13,15,31]. Consequently, it may warrant specific control measures where it exists, particularly when causing a nuisance biting issue to local communities [22]. Following high levels of nuisance biting of people at the index site over many years [22], targeted biocidal larviciding by local pest control operators and subsequent land management undertaken by the landowner in conjunction with a local conservation organisation in early 2025 has modified much of the habitat to markedly reduce the area of floodplain in an attempt to reduce densities of *Ae. vexans*.

Phylogenetic analysis of the UK WNV detection indicates a close evolutionary relationship to WNV lineage 1a sequences detected in the USA, the Middle East and Europe. However, this is based on a short conserved region of the WNV genome and, consequently, we are unable to determine the geographic origin of this incursion [32].

Whilst WNV incursion into the UK is most likely to occur through either long- or short-distance bird migration from mainland Europe [12], natural-or human-facilitated mosquito movement cannot be discounted. Human-facilitated mosquito movement has been implicated as a possible incursion method of WNV into North America in 1999 [33], and has been highlighted as a means of incursion into other island ecosystems [34]. Although the origin of this incursion remains uncertain, current climatic conditions influencing the extrinsic incubation period suggest that WNV would not be able to circulate sustainably in the UK at the index site, [35].

Whilst our detection of WNV RNA in a potential mosquito bridge vector indicates that the UK is permissive for incursion events, the public health risk of WNV is currently considered very low [36]. An integrated approach to surveillance, involving vectors, wildlife and domestic animals, is essential to rapidly identify the temporo-spatial presence of infection and to mitigate transmission to humans.

## Acknowledgements

The authors would like to thank the funders of this project (UKRI and DEFRA) for their ongoing financial support. Additionally, Holly Coombes, Ben Clifton and Ben Mollet for their assistance with ONT GridION and the Central Sequencing Unit (Animal and Plant Health Agency). We would also like to thank staff and volunteers at bird observatories and in nature reserves who have run mosquito traps for the project.

## Notes

**Funding statement** This research was funded by the UK Research and Innovation (UKRI) and Department for Environment, Food and Rural Affairs (Defra) through Vector-Borne RADAR (Real-time Arbovirus Detection And Response) (grant no: BB/X017990/1). While additional funding was provided by DEFRA and the Scottish and Welsh Governments through project SV3045. AAC and BL were funded by Research England.

**Conflict of interest** The authors declare no competing interests.

### Competing Interest Statement

The authors have declared no competing interest.

## References

1. Campbell GL, Marfin AA, Lanciotti RS, Gubler DJ. West Nile virus. Lancet Infect Dis [Internet]. 2002 Sep;2(9):519–29.

2. Zeller HG, Schuffenecker I. West Nile Virus: An Overview of Its Spread in Europe and the Mediterranean Basin in Contrast to Its Spread in the Americas. Eur J Clin Microbiol Infect Dis [Internet]. 2004 Mar 1;23(3):147–56.

3. Sikkema RS, Schrama M, van den Berg T, Morren J, Munger E, Krol L, et al. Detection of West Nile virus in a common whitethroat (Curruca communis) and Culex mosquitoes in the Netherlands, 2020. Euro Surveill [Internet]. 2020 Oct;25(40):2001704.

4. Ziegler U, Bergmann F, Fischer D, Müller K, Holicki CM, Sadeghi B, et al. Spread of West Nile Virus and Usutu Virus in the German Bird Population, 2019–2020. Microorganisms [Internet]. 2022 Apr 12;10(4):807.

5. Heus P, Kolodziejek J, Camp J V, Dimmel K, Bagó Z, Hubálek Z, et al. Emergence of West Nile virus lineage 2 in Europe: Characteristics of the first seven cases of West Nile neuroinvasive disease in horses in Austria. Transbound Emerg Dis [Internet]. 2020 May 24;67(3):1189–97.

6. Young JJ, Haussig JM, Aberle SW, Pervanidou D, Riccardo F, Sekulić N, et al. Epidemiology of human West Nile virus infections in the European Union and European Union enlargement countries, 2010 to 2018. Euro Surveill [Internet]. 2021 May;26(19):2001095.

7. García San Miguel Rodríguez-Alarcón L, Fernández-Martínez B, Sierra Moros MJ, Vázquez A, Julián Pachés P, García Villacieros E, et al. Unprecedented increase of West Nile virus neuroinvasive disease, Spain, summer 2020. Eurosurveillance [Internet]. 2021 May 13;26(19):2002010.

8. Schopf F, Sadeghi B, Bergmann F, Fischer D, Rahner R, Müller K, et al. Circulation of West Nile virus and Usutu virus in birds in Germany, 2021 and 2022. Infect Dis (Auckl) [Internet]. 2025 Mar 4;57(3):256–77.

9. Münger E, Atama N, van Irsel J, Blom R, Krol L, van den Berg TJ, et al. Emergence and Dynamics of Usutu and West Nile Viruses in the Netherlands, 2016-2022 [Internet]. bioRxiv. Cold Spring Harbor Laboratory; 2024. p. 2012–24.

10. Wildlife: GB disease surveillance and emerging threats reports [Internet]. 2024. Available from: https://www.gov.uk/government/publications/wildlife-gb-disease-surveillance-and-emerging-threats-reports

11. Folly AJ, Lawson B, Lean FZ, McCracken F, Spiro S, John SK, et al. Detection of Usutu virus infection in wild birds in the United Kingdom, 2020. Euro Surveill [Internet]. 2020 Oct;25(41):2001732.

12. Bessell PR, Robinson RA, Golding N, Searle KR, Handel IG, Boden LA, et al. Quantifying the Risk of Introduction of West Nile Virus into Great Britain by Migrating Passerine Birds. Transbound Emerg Dis [Internet]. 2016 Oct;63(5):e347–59.

13. Medlock JM, Cull B, Vaux AGC, Irwin AG. The mosquito Aedes vexans in England. Vet Rec [Internet]. 2017 Sep 2;181(9):243–243.

14. Abbott AJ, Gardner BL, Wilson R, Biddlecombe SM, Vaux AGC, Medlock JM. Update on Aedes vexans distribution in the United Kingdom. Vet Rec. 2025; Forethcoming.

15. Börstler J, Jöst H, Garms R, Krüger A, Tannich E, Becker N, et al. Host-feeding patterns of mosquito species in Germany. Parasit Vectors [Internet]. 2016 Dec 3;9(1):318.

16. Biteye B, Fall AG, Seck MT, Ciss M, Diop M, Gimonneau G. Host-feeding patterns of Aedes (Aedimorphus) vexans arabiensis, a Rift Valley Fever virus vector in the Ferlo pastoral ecosystem of Senegal. Samy AM, editor. PLoS One [Internet]. 2019 Oct 4;14(10):e0215194.

17. Tiawsirisup S, Kinley JR, Tucker BJ, Evans RB, Rowley WA, Platt KB. Vector Competence of Aedes vexans (Diptera: Culicidae) for West Nile Virus and Potential as an Enzootic Vector. J Med Entomol [Internet]. 2008 May 1;45(3):452–7.

18. Gendernalik A, Weger-Lucarelli J, Garcia Luna SM, Fauver JR, Rückert C, Murrieta RA, et al. American Aedes vexans Mosquitoes are Competent Vectors of Zika Virus. Am J Trop Med Hyg [Internet]. 2017 Jun;96(6):1338–40.

19. Mravcová K, Camp J V, Hubálek Z, Šikutová S, Vaux AGC, Medlock JM, et al. Ťahyňa virus-A widespread, but neglected mosquito-borne virus in Europe. Zoonoses Public Health [Internet]. 2023 Aug;70(5):371–82.

20. Ndiaye EH, Fall G, Gaye A, Bob NS, Talla C, Diagne CT, et al. Vector competence of Aedes vexans (Meigen), Culex poicilipes (Theobald) and Cx. quinquefasciatus Say from Senegal for West and East African lineages of Rift Valley fever virus. Parasit Vectors [Internet]. 2016 Dec 20;9(1):94.

21. Armstrong PM, Andreadis TG. Eastern Equine Encephalitis Virus in Mosquitoes and Their Role as Bridge Vectors. Emerg Infect Dis [Internet]. 2010 Dec;16(12):1869–74.

22. Vaux AGC, Watts D, Findlay-Wilson S, Johnston C, Dallimore T, Drage P, et al. Identification, surveillance and management of Aedes vexans in a flooded river valley in Nottinghamshire, United Kingdom. J Eur Mosq Control Assoc [Internet]. 2021 Nov 23;39(1):15–25.

23. Linke S, Ellerbrok H, Niedrig M, Nitsche A, Pauli G. Detection of West Nile virus lineages 1 and 2 by real-time PCR. J Virol Methods [Internet]. 2007 Dec;146(1– 2):355–8.

24. Eiden M, Vina-Rodriguez A, Hoffmann B, Ziegler U, Groschup MH. Two new real-time quantitative reverse transcription polymerase chain reaction assays with unique target sites for the specific and sensitive detection of lineages 1 and 2 West Nile virus strains. J Vet Diagn Invest [Internet]. 2010 Sep;22(5):748–53.

25. Johnson N, Wakeley PR, Mansfield KL, McCracken F, Haxton B, Phipps LP, et al. Assessment of a Novel Real-Time Pan-Flavivirus RT-Polymerase Chain Reaction. Vector-Borne Zoonotic Dis [Internet]. 2010 Sep;10(7):665–71.

26. Li H, Durbin R. Fast and accurate short read alignment with Burrows–Wheeler transform. Bioinformatics [Internet]. 2009 Jul 15;25(14):1754–60.

27. Li H, Handsaker B, Wysoker A, Fennell T, Ruan J, Homer N, et al. The Sequence Alignment/Map format and SAMtools. Bioinformatics [Internet]. 2009 Aug 15;25(16):2078–9.

28. Danecek P, Bonfield JK, Liddle J, Marshall J, Ohan V, Pollard MO, et al. Twelve years of SAMtools and BCFtools. Gigascience [Internet]. 2021 Jan 29;10(2):giab008. 2

29. Koch RT, Erazo D, Folly AJ, Johnson N, Dellicour S, Grubaugh ND, et al. Genomic epidemiology of West Nile virus in Europe. One Heal [Internet]. 2024 Jun;18:100664.

30. Davis E, Velez J, Hamik J, Fitzpatrick K, Haley J, Eschliman J, et al. Evidence of Lineage 1 and 3 West Nile Virus in Person with Neuroinvasive Disease, Nebraska, USA, 2023. Emerg Infect Dis [Internet]. 2024 Oct;30(10):2090–8.

31. Molaei G, Andreadis TG. Identification of avian-and mammalian-derived bloodmeals in Aedes vexans and Culiseta melanura (Diptera: Culicidae) and its implication for West Nile virus transmission in Connecticut, USA. J Med Entomol. 2006;43(5):1088– 93.

32. Banet-Noach C, Malkinson M, Brill A, Samina I, Yadin H, Weisman Y, et al. Phylogenetic relationships of West Nile viruses isolated from birds and horses in Israel from 1997 to 2001. Virus Genes. 2003;26:135–41.

33. Lanciotti RS, Roehrig JT, Deubel V, Smith J, Parker M, Steele K, et al. Origin of the West Nile virus responsible for an outbreak of encephalitis in the northeastern United States. Science (80-). 1999;286(5448):2333–7.

34. Kilpatrick AM, Daszak P, Goodman SJ, Rogg H, Kramer LD, Cedeno V, et al. Predicting Pathogen Introduction: West Nile Virus Spread to Galápagos. Conserv Biol [Internet]. 2006 Aug 24;20(4):1224–31.

35. Ewing DA, Purse B V, Cobbold CA, White SM. A novel approach for predicting risk of vector-borne disease establishment in marginal temperate environments under climate change: West Nile virus in the UK. J R Soc Interface [Internet]. 2021 May 26;18(178):20210049.

36. Qualitative assessment of the risk that West Nile virus presents to the UK human health population [Internet]. 2020.

37. Vaux AGC, Abbott AJ, Johnston CJ, Hawkes FM, Hopkins RJ, Cull B, et al. An update on the ecology, seasonality and distribution of Culex modestus in England. J Eur Mosq Control Assoc [Internet]. 2024 Apr 19;42(2):77–95.

38. Quick J, Grubaugh ND, Pullan ST, Claro IM, Smith AD, Gangavarapu K, et al. Multiplex PCR method for MinION and Illumina sequencing of Zika and other virus genomes directly from clinical samples. Nat Protoc [Internet]. 2017 Jun 24;12(6):1261–76.

