## Supplementary Information for "Detection of West Nile Virus via Retrospective Mosquito Arbovirus Surveillance in the United Kingdom"

**Supplementary Methodology**

**Mosquito sampling at Gamston**

Of the 31,582 mosquitoes collected and screened for WNV RNA as part of the Vector-Borne RADAR project, the 2,000 female *Ae. vexans* from Gamston were selected due to their vectoral-capacity for Tahyna virus [19]. Due to laboratory capacity restraints, it was not possible to process the full collection of 44,080 mosquitoes (of which 99.99% were *Ae.vexans*) from Gamston in this project. Whilst additional *Ae. vexans* samples were collected from a second site in Suffolk 2024 (Supplementary Table 1), it was not possible to collect samples at Gamston due to construction works (see main text).

The authors note that the primary vectors of WNV (specifically *Cx. pipiens s.s.* and *Cx. modestus*) were not trapped at this site in 2023 for two reasons. Firstly, the octenol lure used during trapping is specific for mammalophagic species and thus would not attract the ornithophagic *Cx. pipiens pipiens.* Secondly, *Cx. modestus* and *Cx. p. molestus* are only found in localised areas of the UK and have not been recorded at the Gamston site to date [37].

**Nucleotide extraction**

Mosquito pools were homogenised in 500 µl of MagMAX^TM^ CORE Lysis Solution (Thermo Fisher, UK), using a 5 mm steel bead and a TissueLyser2 (Thermo Fisher, UK) for 3 minutes at 30 rps. Initial nucleotide extraction was then conducted using 350 µl of this lysate and a KingFisher Flex Purification System (Thermo Fisher, UK) following the manufacturer’s instructions and eluted in 50 µl of MagMAX^TM^ CORE elusion buffer (Thermo Fisher, UK).

Where WNV RNA was detected, viral RNA was re-extracted from 150 µl lysate utilising a QIAamp Viral RNA Extraction mini kit (QIAGEN, UK) following the manufacturer’s instructions and eluted in 50 µl of Buffer AVL (QIAGEN, UK).

**WNV Multiplex PCR**

Positive sample pools were subjected to a multiplex PCR to amplify consecutive 400 bp regions of the WNV lineage 1a genome. Reverse transcription was conducted on viral RNA extracts using LunaScript© (New England Biolabs, USA) according to the manufacturer’s instructions. Primers obtained from PrimalScheme (yale-west-nile-virus/400/v1.0.0) [38], were separated into two pools (as indicated in Supplementary Table 3). PCR reactions comprised 12.5 µl Q5® High-Fidelity 2x master mix (New England Biolabs, USA), 4 µl of respective primer pool (1 or 2), 2.5 µl of synthesised cDNA and 6 µl of molecular grade water.

Reactions were subjected to conventional PCR using the following cycling conditions: initial denaturation at 98 ºC for 30 seconds; 35 cycles of 98 ºC for 10 seconds, 62 ºC for 30 seconds and 72 ºC for 30 seconds, and a final 5-minute extension period at 72 ºC.

|  | March-October 2023 | | January-November 2024 | | January-March 2025 | |  | |
| --- | --- | --- | --- | --- | --- | --- | --- | --- |
| Mosquito species | Total Female (*n*) | Total Male (*n*) | Total Female (*n*) | Total Male (*n*) | Total Female (*n*) | Total Male (*n*) | Total individuals (*n*) | Total number of pools (*n*) |
| *Aedes cantans/annulipes* | 901 | 1 | 1169 | 0 | 0 | 0 | 2071 | 252 |
| *Aedes caspius* | 250 | 0 | 86 | 0 | 0 | 0 | 336 | 48 |
| *Aedes cinereus/geminus* | 102 | 0 | 2298 | 0 | 0 | 0 | 2400 | 260 |
| *Aedes communis* | 0 | 0 | 8 | 0 | 0 | 0 | 8 | 3 |
| *Aedes detritus* | 1854 | 0 | 1011 | 0 | 0 | 0 | 2865 | 323 |
| *Aedes dorsalis* | 0 | 0 | 3 | 0 | 0 | 0 | 3 | 2 |
| *Aedes flavescens* | 374 | 0 | 59 | 0 | 0 | 0 | 433 | 49 |
| *Aedes geniculatus* | 4 | 1 | 1 | 0 | 0 | 0 | 6 | 4 |
| *Aedes leucomelas* | 0 | 0 | 1 | 0 | 0 | 0 | 1 | 1 |
| *Aedes punctor* | 0 | 0 | 4 | 0 | 0 | 0 | 4 | 3 |
| *Aedes rusticus* | 673 | 0 | 51 | 0 | 0 | 0 | 724 | 89 |
| *Aedes sticticus* | 0 | 0 | 2 | 0 | 0 | 0 | 2 | 2 |
| *Aedes vexans* | **2000** | 0 | 1380 | 0 | 0 | 0 | 3380 | 338 |
| *Aedes spp.** | 1 | 0 | 12 | 0 | 0 | 0 | 13 | 4 |
| *Anopheles claviger* | 3229 | 15 | 815 | 1 | 0 | 0 | 4060 | 453 |
| *Anopheles maculipennis s.l.* | 51 | 4 | 44 | 1 | 0 | 0 | 100 | 34 |
| *Anopheles plumbeus* | 58 | 0 | 89 | 0 | 0 | 0 | 147 | 38 |
| *Coquillettidia richardii* | 884 | 0 | 1438 | 0 | 0 | 0 | 2322 | 285 |
| *Culex modestus* | 1026 | 0 | 1907 | 0 | 0 | 0 | 2933 | 311 |
| *Culex pipiens s.l.* | 6522 | 7 | 949 | 2 | 514 | 0 | 7994 | 859 |
| *Culex spp.** | 14 | 0 | 27 | 0 | 0 | 0 | 41 | 8 |
| *Culiseta annulata* | 949 | 10 | 318 | 0 | 5 | 0 | 1282 | 206 |
| *Culiseta morsitans* | 435 | 0 | 16 | 0 | 0 | 0 | 451 | 65 |
| Unknown | 0 | 0 | 6 | 0 | 0 | 0 | 6 | 2 |
| Total | 19327 | 38 | 11694 | 4 | 519 | 0 | 31582 | 3639 |

Supplementary Table 1: The total number (*n*) of individual mosquitoes caught between March 2023-March 2025, inclusive, and analysed as part of the Vector-Borne RADAR project. Samples were pooled (≤13) by species, sex, location and collection date. All samples tested negative for WNV RNA, except two pools of the 2000 *Aedes vexans* (marked in bold). Where possible mosquito specimens were kept at -20 ºC immediately following capture, all samples were transferred to -80 ºC for long-term storage. * indicates specimens identified to genus level but unidentifiable at species level.

**Supplementary Table 2**. The sixty-five West Nile virus sequences obtained from GenBank used in phylogenetic analysis. Sequences were selected from the available Genbank repository with a broad temporal and spatial representation (six from Africa (1958-2006); two from Asia (1998-2005); 18 from Europe (2916-2018) and 38 from North America (1999-2022) to provide coverage of the West Nile virus evolutionary history.

| **Accession number** | **Date** | **Location Country** | **Continent** | **Lineage** |
| --- | --- | --- | --- | --- |
| HM147824 | 1958 | Democratic Republic of Congo | Africa | L2 |
| HM147822 | 1958 | South Africa | Africa | L2 |
| HM147823 | 1988 | Madagascar | Africa | L2 |
| OP870459 | 1998 | Senegal | Africa | L1a |
| OP870461 | 2006 | Senegal | Africa | L1a |
| ON813219 | 2006 | Senegal | Africa | L2 |
| AF481864 | 1998 | Israel | Asia | L1a |
| AY688948 | 2005 | Israel | Asia | L2 |
| OM302316 | 2016 | Spain | Europe | L1a |
| OM302319 | 2017 | Spain | Europe | L1a |
| OM302317 | 2018 | Spain | Europe | L1a |
| MH021189 | 2018 | Belgium | Europe | L2 |
| MH924836 | 2018 | Germany | Europe | L2 |
| OP713602 | 2020 | Spain | Europe | L1a |
| OP713603 | 2020 | Spain | Europe | L1a |
| MW036634 | 2020 | Netherlands | Europe | L2 |
| OP762597 | 2020 | Netherlands | Europe | L2 |
| MW036633 | 2020 | Netherlands | Europe | L2 |
| MW036634 | 2020 | Netherlands | Europe | L2 |
| OP609793 | 2022 | Italy | Europe | L1a |
| OP609798 | 2022 | Italy | Europe | L1a |
| OP609813 | 2022 | Italy | Europe | L1a |
| PQ450190 | 2024 | Poland | Europe | L2 |
| PQ483500 | 2024 | Spain | Europe | L2 |
| PQ481864 | 2024 | Spain | Europe | L2 |
| MN619803 | 2018 | Russia | Europe | L2 |
| NC009942 | 1999 | New York, USA | North America | L1a |
| AF404756 | 2000 | New York, USA | North America | L1a |
| KJ786934 | 2001 | New York, USA | North America | L1a |
| HQ671697 | 2001 | USA | North America | L1a |
| KX547194 | 2002 | New York, USA | North America | L1a |
| KX547287 | 2003 | New York, USA | North America | L1a |
| JQ700437 | 2003 | Ney York, USA | North America | L1a |
| HM488145 | 2004 | USA | North America | L1a |
| MH819448 | 2005 | Canada | North America | L1a |
| OQ721162 | 2005 | Minnesota, USA | North America | L1a |
| JF415914 | 2005 | Texas, USA | North America | L1a |
| MH819469 | 2006 | Canada | North America | L1a |
| JF415916 | 2006 | Texas, USA | North America | L1a |
| JF415915 | 2006 | Texas, USA | North America | L1a |
| JF957171 | 2007 | Indiana, USA | North America | L1a |
| JF415918 | 2007 | Texas, USA | North America | L1a |
| JF957170 | 2007 | USA | North America | L1a |
| JF957173 | 2008 | Arizona, USA | North America | L1a |
| JN183886 | 2008 | New York, USA | North America | L1a |
| JF415927 | 2008 | Texas, USA | North America | L1a |
| JF957176 | 2009 | Arizona, USA | North America | L1a |
| OQ721116 | 2009 | California, USA | North America | L1a |
| JF488095 | 2009 | New York, USA | North America | L1a |
| JF957183 | 2009 | Texas, USA | North America | L1a |
| JQ700438 | 2011 | USA | North America | L1a |
| KY216149 | 2012 | New York, USA | North America | L1a |
| MH819447 | 2013 | Canada | North America | L1a |
| OK631659 | 2013 | New York, USA | North America | L1a |
| MH819446 | 2015 | Canada | North America | L1a |
| OK631660 | 2015 | New York, USA | North America | L1a |
| MT967990 | 2015 | New York, USA | North America | L1a |
| OQ721131 | 2016 | California, USA | North America | L1a |
| MT967996 | 2017 | New York, USA | North America | L1a |
| OQ721122 | 2018 | Illinois, USA | North America | L1a |
| MT968010 | 2018 | New York, USA | North America | L1a |
| OQ721111 | 2018 | Ohio, USA | North America | L1a |
| MZ595325 | 2021 | New York, USA | North America | L1a |
| MZ595324 | 2022 | New York, USA | North America | L1a |

Supplementary Table 3: Primers used in the multiplex primer tiling PCR, available at PrimalScheme [38]. Pools 1 and 2 were run in separate PCR reactions and the products combined prior to barcoding during GridION sample preparation.

| **Forward** | **Reverse** | **Pool** |
| --- | --- | --- |
| GCCTGTGTGAGCTGACAAACTT | GATAGCACTGGTCAAGGTCCCT | 1 |
| GCGTTCTTCAGGTTCACAGCAA | GACTTTGTGCACCAACAGTCGA | 2 |
| TGTCAGAGCAATGGATGTGGGA | CTGTTGCTCATTCCAAGGCAGT | 1 |
| TCATTGGTTGGATGCTTGGGAG | TGTGTCAATGCTTCCTTTGCCA | 2 |
| AATGACAAACGTGCTGACCCAG | CACTCACGATGGACCAAGAACG | 1 |
| ATATGGAGAGGTGACAGTGGACT | AAAGCCTTTGAACAGACGCCAT | 2 |
| TTCAAGCAACACTGTCAAGTTAAC | CTGCTTCCAGACTTGTGCCAAT | 1 |
| GCCAACGCTAAGGTCCTGATTG | ACGGAGAGGAAGAGCAGAACTC | 2 |
| GCATGTCCTGGATAACGCAAGG | AACCACGACACTAAGGTCCACA | 1 |
| TCTACGATCAGTTTCCAGACTGGA | GTTGTTCTTGACAGCCGTTCCA | 2 |
| TGGAGGATTTTGGATTTGGTCTCAC | AGTCAATCTCTACCCGGCCTTC | 1 |
| GGCGATGGAATCCTTGAGAGTG | GGCCCAACTGAAAAGGGTCAAT | 2 |
| CGGCTGTTGGTATGGTATGGAG | TTCTCCTGGTTGGTCCATCTCG | 1 |
| CTAATTCGGGAGGAGACGTGGT | CCTGATCAAGCTGCCTATTCCG | 2 |
| ATCAAACGTGGTTGTTCCGCTG | CACGAAAGCAGCAAACATGAGC | 1 |
| TGTCGGCCTGATGTTTGCCATC | GGTCGTGTCCCCCTTTTTGTAC | 2 |
| TGCCCTCAGTAGTTGGATTTTGG | TTTGAACACCCCTGGTTTCGTC | 1 |
| TGGAAATTGCAGCACAAGTGGA | GGCCTCTTTGATGATCTGTGGC | 2 |
| CGGATTCGAACCTGAGATGCTG | AACCTCTTGCTGCAATGCTAGC | 1 |
| ACAGGCTGATGTCTCCTCACAG | ACATTTTGGGTACTCCGTCTCGT | 2 |
| GTGCCTAGTGTCAAGATGGGGA | GATTCGTGCCTCAGTCCAATGG | 1 |
| CGGTAGAAATCCGTCGCAAGTT | GCGGCCTCAGAATCTTCCTTTC | 2 |
| AGGTGGTGCTTTGATGGTCCTA | ACTCCCATGGTCATCACACTCA | 1 |
| TGCAGAGAAAGGAGGAAGAGCT | CCTTGACCTCAATTCTTTGCCCA | 2 |
| TTGTGTCATGACCCTTGTGAGC | GGAACCATGTAGGCATAGTGGC | 1 |
| CTTCGTCGATGTTGGAGTGTCG | CTCCATTCTCCCAAAGCGTCAC | 2 |
| TCTTGGTGTCTCTAGCTGCAGT | CAGTTTTGCTGTGCCCCTAGAG | 1 |
| GTACCGCAAAGAGGCCATCATC | TTGACGAGGACTCTCCGATGTC | 2 |
| TGGAACATTGTCACCATGAAGAGT | CTTCCTCGTATTGGGGTCCCTT | 1 |
| CGAGCTTCAGGCAATGTGGTAC | AGGGAGTAGTGTCAGTCATGGC | 2 |
| CAGTTCGCTGGTCAATGGAGTG | AATGCAAGTGTGACATTCCCCC | 1 |
| GAGCAGAATCAATGGAGGAGCG | TCCAAGTCAGCTCTCGTGATGC | 2 |
| TTACATCCTGCGTGAAGTTGGC | TCTCCACATCATCTGGGCCAAT | 1 |
| TGTCACCTACGCCCTAAACACT | GGAGATATGCGAGCTCTGCCTA | 2 |
| CAGGTTCCATTTTGCTCAAACCA | TCACTCCATTTCTCCACTGGGG | 1 |
| CATGCAGGAGGAGAGTGGATGA | CGGCCTGACTTTCTCCTCAAAA | 2 |
| TGGTTGAGGACACAGTACTGTAGA | GTCCTTTCGCCCTGGTTAACAC | 1 |
| GCGAAGTGATCCATGTAAGCCT | TGTGCAGAGCAGAAGATCTCCT | 2 |
